# Speech Synthesis from Electrocorticography during Imagined Speech Using a Transformer-Based Decoder and a Pretrained Vocoder

**DOI:** 10.1101/2024.08.21.608927

**Authors:** Shuji Komeiji, Kai Shigemi, Takumi Mitsuhashi, Yasushi Iimura, Hiroharu Suzuki, Hidenori Sugano, Koichi Shinoda, Kohei Yatabe, Toshihisa Tanaka

## Abstract

Synthesizing speech from Electrocorticogram (ECoG) signals recorded during imagined speech remains a challenge due to the absence of synchronized audio signals for training. To address this, we propose a training framework that utilizes audio recorded during overt speech tasks as a surrogate ground truth for imagined speech signals, based on the consistency of the linguistic content. We employed a Transformer-based decoder to generate log-mel spectrograms from imagined speech ECoG, which were then converted into waveform audio using a pre-trained Parallel WaveGAN. In experiments involving ECoG recordings from 13 participants, the synthesized speech achieved dynamic time warping-aligned Pearson correlation coefficients ranging from 0.74 to 0.84 with the proxy targets. These results demonstrate that overt speech audio can serve as an effective training target for reconstructing imagined speech, offering a viable solution for training decoders in the absence of behavioral output.

## 1. Introduction

Brain–Computer Interfaces (BCIs) are technologies that establish a direct pathway between the brain and the external environment. The development of techniques to interpret meanings and messages from recorded brain activity is of paramount importance in BCI research. Generally, the methods for measuring, processing, and analyzing brain activity vary depending on the type of information to be extracted. These targets may include motor actions or motor intentions [1, 2, 3], internal speech [4, 5, 6], or psychological states [7, 8, 9]. In particular, speech-related BCIs (speech BCIs) hold great promise as assistive technologies for patients with lost or significantly impaired speech function due to brain disorders such as aphasia [10], stroke, or amyotrophic lateral sclerosis [11]. These technologies have the potential to significantly enhance communication capabilities for individuals affected by such conditions. To achieve such a BCI, one promising approach is to decode speech signals from invasive electrocorticography (ECoG) recordings acquired via implanted subdural grid electrodes. Compared to surface electroencephalography (EEG), ECoG offers superior spatiotemporal resolution and signal-to-noise ratio, making it particularly suitable for analyzing speech-related brain activity in the high gamma band [12, 13].

Previous studies indicate that speech estimation from ECoG or stereo-EEG (sEEG) signals primarily falls into two categories: textual output [14, 15, 16] and synthesized speech [17, 18, 19]. Most studies producing synthesized speech in audio format adopt a two-stage approach: the first stage involves estimating a speech representation, followed by the synthesis of audio signals from that representation. Herff et al. [17] directly estimated discrete audio units from ECoG signals corresponding to the participant’s voice and concatenated them to generate continuous speech. Anumanchipalli et al. [18] employed a bidirectional long short-term memory (BLSTM) network [20] to decode articulatory kinematics as an intermediate speech representation from cortical activity, which was subsequently transformed into audio signals. Angrick et al. [19] estimated log-mel spectrograms by applying linear discriminant analysis to predict energy levels across spectral bins. Then, these spectrograms were converted into audio using the Griffin–Lim algorithm [21].

The quality of synthesized speech has significantly improved with the advent of neural vocoders such as WaveNet [22], WaveGlow [23], and Parallel WaveGAN [24]. Angrick et al. [25] used WaveNet to reconstruct spoken audio from ECoG signals. They first estimated log-mel spectrograms using DenseNets [26], which were then input to WaveNet. Kohler et al. [27] utilized WaveGlow for spectrogram-to-speech conversion, estimating log-mel spectrograms from cortical activity using a gated recurrent unit (GRU) [28], which captured temporal dependencies more effectively than DenseNets. Shigemi et al. [29] employed a Transformer-based model [30] to estimate log-mel spectrograms from ECoG signals recorded during overt (vocalized) speech tasks and used a pre-trained Parallel WaveGAN [24] to synthesize high-quality speech from the estimated spectrograms.

Previous studies have focused on synthesizing speech from neural activity during overt speech. Still, the ultimate goal of speech BCIs is to decode or synthesize covert (imagined) speech. However, collecting covert speech data for training poses a fundamental challenge: the internal state of covert speech cannot be observed externally, rendering ground-truth audio signals unavailable for supervised training. To address this challenge, we developed a task presentation paradigm termed “Karaoke-like text highlighting” and demonstrated the feasibility of decoding text from covert speech using a Transformer model trained on overt speech data collected via this paradigm [31]. These findings suggest that covert speech may share common neural signal patterns with overt speech, an idea supported by several previous studies [32, 5, 33, 34]. Based on these shared neural mechanisms, we hypothesize that a decoding framework trained to reconstruct the participant’s own voice from overt speech neural activity can be directly generalized to synthesize covert speech. This approach, which extends the overt speech decoding synthesis by Shigemi et al. [29], enables us to synthesize covert speech by utilizing the paired audio signals from overt speech tasks, thereby circumventing the lack of acoustic training data for covert speech.

To validate this hypothesis, we utilized ECoG recordings from 13 participants, identical to those used in our previous study [31]. These signals were used to synthesize speech by combining either a BLSTM-based or a Transformer-based decoder with a pre-trained Parallel WaveGAN. The quality and accuracy of the synthesized speech derived from both overt and covert speech were evaluated using two metrics: a Pearson Correlation Coefficient (PCC) calculated after temporal alignment via a dynamic time warping (DTW) algorithm [35] and a token error rate (TER) from dictation test results. The results demonstrated that covert speech could be synthesized with high fidelity (PCC of 0.74 to 0.84) and semantic accuracy, as evidenced by a TER of 47.2%, which is significantly below the 51.0% when Gaussian noise was input instead of actual ECoG signals (*p <* 0.01).

## 2. Methods

### 2.1 Participants

In this study, the 13 volunteer participants (labeled js4 to js16; see Table 1) were undergoing treatment for drug-resistant temporal lobe epilepsy at the Department of Neurosurgery, Juntendo University Hospital. The group comprised seven male and six female individuals, with a mean age of 26.1 (range: 8–41) years. ECoG electrode arrays were surgically implanted on the cortical surface of each participant to localize seizure foci. Prior to implantation, the language-dominant hemisphere was identified using functional magnetic resonance imaging (fMRI). ECoG and audio data were collected from all 13 participants, each of whom provided informed consent. The experimental protocol for data acquisition was approved by the Medical Research Ethics Committee of Juntendo University Hospital, and the data analysis was approved by the Ethics Committee of the Tokyo University of Agriculture and Technology. All research procedures were conducted in accordance with relevant institutional guidelines and regulations.

**Table 1.**
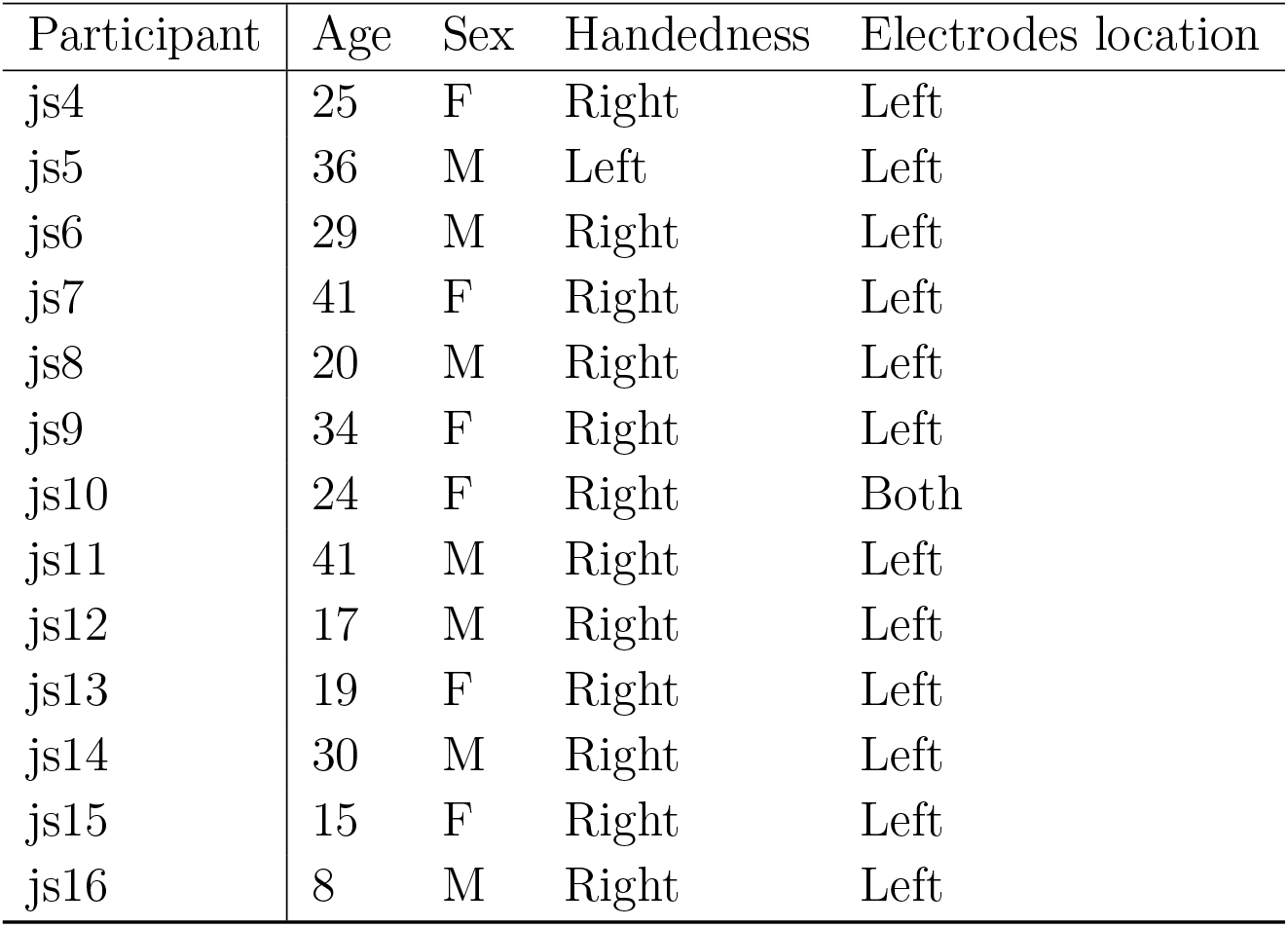
Clinical profiles for 13 participants. The age, sex, handedness, and electrode location are listed.

| Participant | Age | Sex | Handedness | Electrodes location |
| --- | --- | --- | --- | --- |
| js4 | 25 | F | Right | Left |
| js5 | 36 | M | Left | Left |
| js6 | 29 | M | Right | Left |
| js7 | 41 | F | Right | Left |
| js8 | 20 | M | Right | Left |
| js9 | 34 | F | Right | Left |
| js10 | 24 | F | Right | Both |
| js11 | 41 | M | Right | Left |
| js12 | 17 | M | Right | Left |
| js13 | 19 | F | Right | Left |
| js14 | 30 | M | Right | Left |
| js15 | 15 | F | Right | Left |
| js16 | 8 | M | Right | Left |

### 2.2 Listeners for Dictation Test

For the dictation test to evaluate the synthesized speech (see Section 2.13.3 for details), we recruited 14 native Japanese-speaking listeners with normal hearing to the synthesized speech. This test was approved by the Ethics Committee of the Tokyo University of Agriculture and Technology.

### 2.3 Data Acquisition

ECoG signals were recorded using intracranial electrodes implanted on the cortical surface. An epidural electrode was used as the ground (GND). All signals, except for the one from the epidural electrode, were recorded as differential signals with respect to the GND and amplified using a biomedical amplifier (g.HIAMP, g.tec Medical Engineering GmbH, Austria). Electrooculograms were simultaneously recorded using four disposable electrodes to monitor eye movements. In addition to ECoG signals, audio signals, such as participants’ vocalizations, were recorded using a microphone (ECM360, SONY, Japan) and digitized in synchrony with the ECoG recordings. Both ECoG and audio signals were digitized at a sampling rate of 9, 600 Hz.

Trigger signals indicating event status changes were recorded using a photodetector circuit, which combined a linear light sensor (LLS05-A, Nanyang Senba Optical and Electronic Co., Ltd., Nanyang, China) with an operational amplifier (NJM3404AD, New Japan Radio, Tokyo, Japan). The photodetector was positioned in a corner of the monitor screen, where pixel color changes were used to indicate event transitions. All signals were recorded using the Simulink toolbox (MathWorks, Natick, MA, USA), which also controlled the biomedical amplifier.

### 2.4 Experimental Design

We recorded ECoG signals under three conditions: auditory speech perception, overt speech, and covert speech. To collect these data, we designed the task schedule shown in Fig. 1a. According to this schedule, participants performed the perception, overt, and covert tasks sequentially, which we collectively refer to as a “track.” For each track, a single sentence was presented. To ensure a balanced dataset, the experiment was designed such that each of the eight unique sentences appeared exactly ten times across the 80 tracks, presented in a randomized order for each participant. Intervals of more than 6 seconds between tasks were included to prevent time-correlated artifacts. If an error occurred during a trial, that trial was repeated, and thus the total number of trials could exceed 80. As a result, we obtained 80 segments of ECoG and audio signals for each task and participant. During the recordings, participants were seated in a chair at a desk equipped with a monitor, a loudspeaker, a microphone, and a keyboard. The monitor and loudspeaker were controlled in accordance with the task schedule.

**Figure 1.**
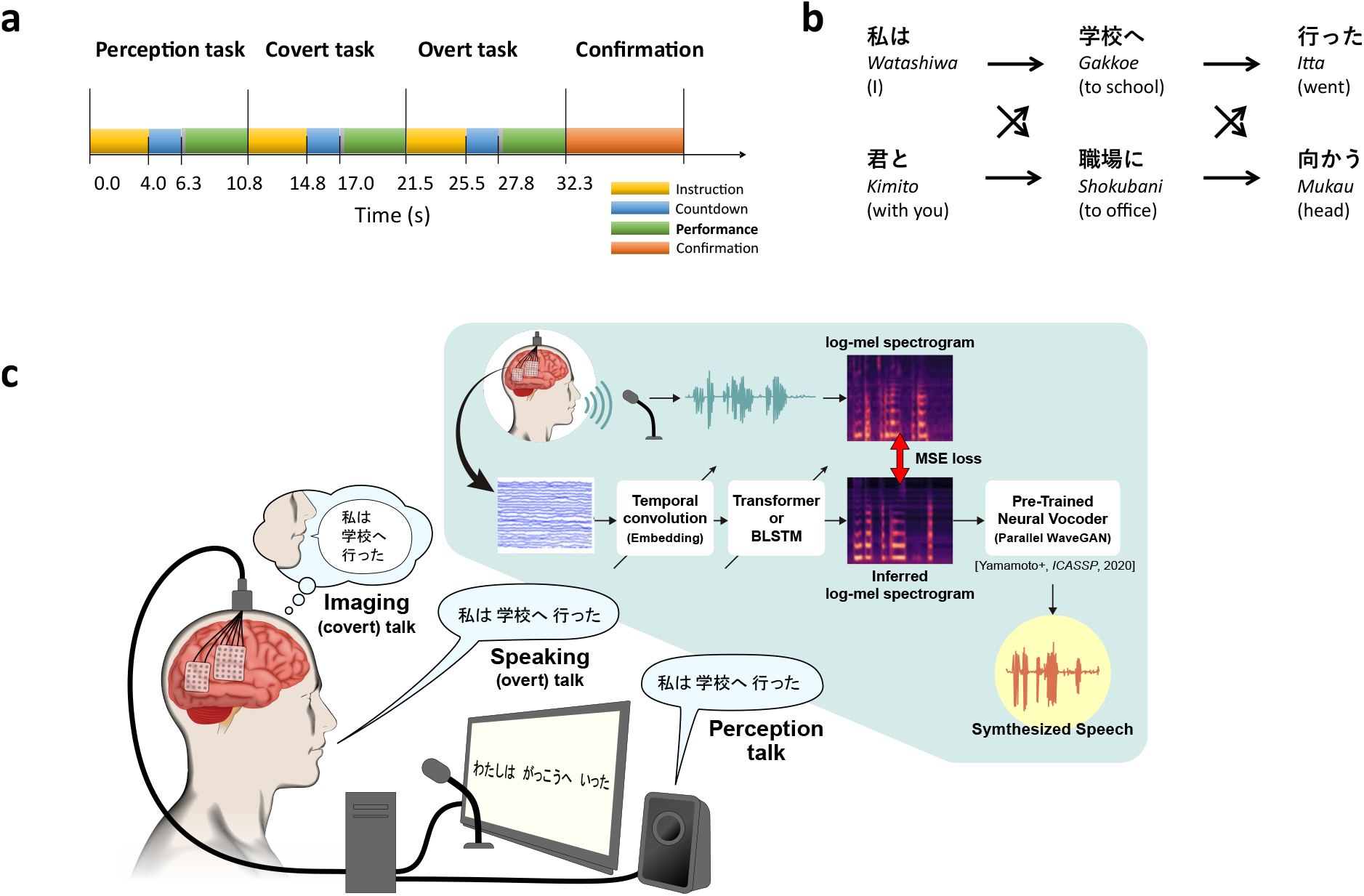
Study overview. **a** Experimental design. A single track lasts over 33 seconds to record sentences for the three tasks. The yellow box indicates task instructions, the blue box indicates a countdown for preparation, and the green box indicates task performance: perception, covert, or overt. The red box indicates the prompt for error judgment. **b** Sentence generation. Sentences consist of three Japanese phrases, each with two options, resulting in a total of eight sentences. The figure depicts the structure of each sentence, showing the Hiragana representation at the top, Romaji (pronunciation) in the middle, and English translation in parentheses below. **c** Decoding process. The process involves a neural network, which is composed of a temporal convolution, a log-mel spectrogram decoder, and a pre-trained neural vocoder.

The track consisted of the perception, overt, and covert tasks in that order. At the beginning of each task, instruction and countdown periods were included to prepare the participants for the upcoming task (yellow and blue boxes depicted in Fig. 1a). In the perception task, participants were instructed to listen to the sentence played from the loudspeaker while gazing at a cross symbol displayed on the monitor. In the covert and overt tasks, participants were asked to read silently or aloud a sentence presented using a “Karaoke-like text highlighting” method on the monitor. For the overt task, participants were instructed to read the highlighted text aloud naturally. For the covert task, they were explicitly instructed to “read the highlighted text silently in your mind as if speaking it, articulating each character at the presented pace, but without producing any vocal sound or physical mouth movement.” Although this paradigm involves visual reading, in this study, we refer to this silent reading task as “covert speech.” While we acknowledge the terminological distinction between spontaneous “imagined speech” and “silent reading,” supervised decoding requires precise temporal alignment between neural activity and target labels. Therefore, we employed this reading-based paradigm to strictly control the timing of the internal speech production, following established protocols in the field (e.g., covert word repetition tasks [13]).

The text highlighting proceeded at a constant rate of 5 characters per second. To prevent participants from reading faster or slower than the highlight speed at the beginning of the sentence, a dummy word—one that does not exist in the Japanese language and carries no meaning but is pronounceable—was inserted at the beginning of the sentence (gray boxes in Fig. 1a). At the end of each track, participants used the keyboard to indicate whether there were any mistakes during the trial (red box in Fig. 1a). If no mistakes were reported, the next track commenced; otherwise, the current track was repeated.

During the experiments, we continuously recorded ECoG signals, an audio signal, and a trigger signal, which indicated the periods of the dummy words and sentences (gray and green boxes in Fig. 1a). The trigger signal was used to trim both the ECoG and audio signals.

### 2.5 Sentences

The sentences displayed to the participants comprised three Japanese phrases (Fig. 1b). The total number of sentences was eight. Therefore, each of the eight unique sentences appeared ten times across the 80 trials per task. Each phrase had two options to generate a sentence. The first phrase was either “watashiwa” (I) or “kimito” (with you), the second phrase was either “gakko:e” (to school) or “shokubani” (to the office), and the third phrase was either “itta” (went) or “mukau” (head). Consequently, we generated eight (= 2 × 2 × 2) sentence patterns.

### 2.6 Decoding Process

We synthesized audio signals from each segment of the ECoG signals by employing the decoding process illustrated in Fig. 1c. The process consists of neural network components, including temporal convolution, a log-mel spectrogram decoder, and a pre-trained neural vocoder.

First, as input to the temporal convolution, the segments of the ECoG signals were preprocessed by extracting envelopes of the high gamma bands and downsampled to a sampling rate of 200 Hz. A detailed explanation of the preprocessing is provided in Section 2.7. Next, the envelopes were fed into the temporal convolution module to effectively downsample the temporal dimension. Then, the output of the temporal convolution was passed to the log-mel spectrogram decoder. The predicted log-mel spectrograms were subsequently converted into speech waveforms using a pretrained neural vocoder.

In this study, we implemented two types of log-mel spectrogram decoders: BLSTM-based and Transformer-based. The architectures of both decoders follow the designs described by Shigemi et al. [29]. A detailed description of the network architecture is provided in Section 2.9.

### 2.7 ECoG Preprocessing

Preprocessing to extract ECoG features from the raw signals for machine learning training and decoding involved the following steps [36]. First, electrodes exhibiting artifacts or excessive noise were excluded based on visual inspection by an epileptologist. The number of electrodes retained for analysis per participant were: 54 for js4, 54 for js5, 54 for js6, 53 for js7, 55 for js8, 71 for js9, 42 for js10, 45 for js11, 39 for js12, 56 for js13, 58 for js14, 65 for js15, and 24 for js16. The spatial coverage of valid electrodes across all participants is illustrated in Fig. 2. Second, we segmented both ECoG and audio signals using trigger timestamps indicating sentence start and end, discarding the initial dummy word periods. The length of each trimmed segment corresponded to the maximum sentence duration recorded during the experiment for each participant (approximately 3.5 s). This ensured uniform input length for the decoder and prevented bias from variable segment durations. Performing trimming prior to further signal processing also mitigated time-correlated artifacts originating from adjacent segments. Third, the trimmed ECoG signals were anti-aliased using a low-pass filter at 200 Hz and downsampled to 400 Hz. Notch filters at 50 and 100 Hz (infinite impulse response) were applied to suppress power line noise and its harmonics. Finally, analytic signal amplitudes were extracted from eight adjacent frequency bands between 70 and 150 Hz. Each band was filtered by a finite impulse response bandpass filter with passbands of 68–78, 74–84, 82–92, 91–102, 101–112, 112–124, 124–136, and 137–150 Hz. These band-specific amplitudes were averaged and downsampled to 200 Hz. Then, the resulting analytic amplitudes were *z*-scored electrode-wise to produce the final ECoG features used for decoding.

**Figure 2.**
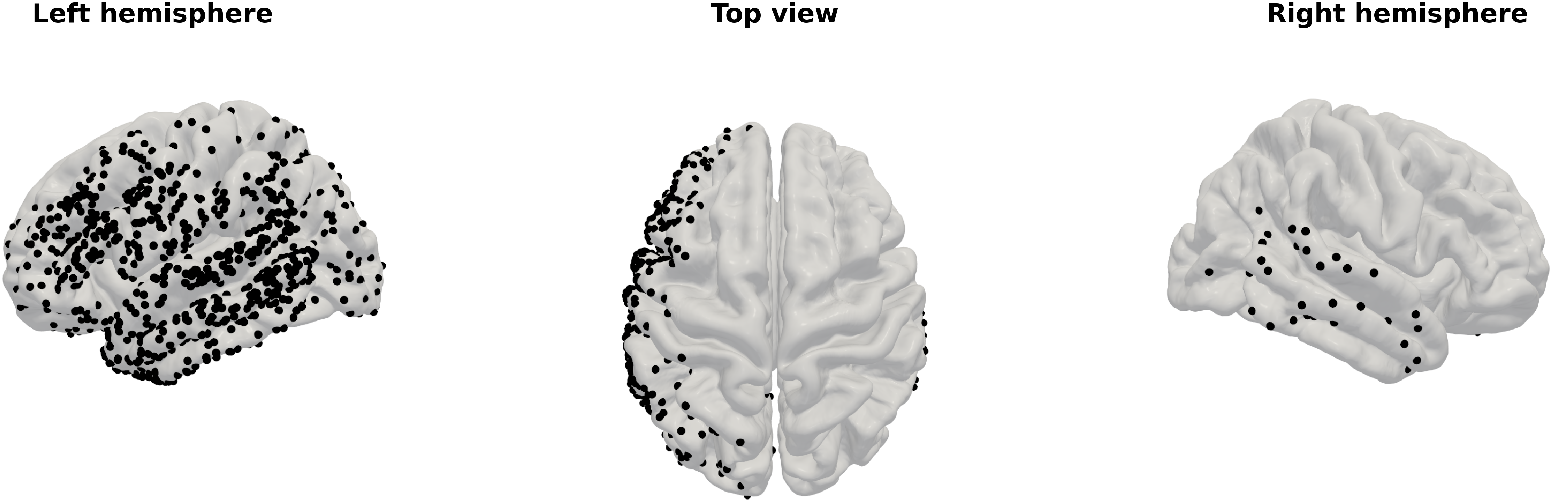
ECoG electrode coverage across all participants.

### 2.8 Audio Preprocessing

The log-mel spectrograms used to train the decoder were computed from the trimmed audio signals using the `Python Speech Features` package [37] following this procedure. First, the audio data were upsampled to 24, 000 Hz. Next, log-mel spectrograms were calculated over the frequency range of 80–7, 600 Hz, which effectively captures the acoustic features relevant for speech [38]. We used 80 mel frequency bins, with a window length of 1, 200 samples and a hop size of 300 samples (corresponding to 12.5 ms). These parameters were chosen to match those of the pre-trained neural vocoder model used for waveform synthesis (see Section 2.10).

### 2.9. Neural Network Architecture‡

We implemented both BLSTM- and Transformer-based architectures for the log-mel spectrogram decoder. BLSTM [39] is a type of recurrent neural network (RNN) designed to efficiently capture both long- and short-term dependencies, making it widely used in ASR and natural language processing (NLP). In contrast, the Transformer [30] employs self-attention mechanisms to model long-range dependencies and has become the state-of-the-art architecture in ASR, NLP, and various other domains such as image processing since its introduction in 2017.

Figure 3 illustrates the detailed network architectures for both decoders. with implementation parameters summarized in Table 2. The decoding process occurs in two stages: a temporal convolutional network and a decoder network.

**Table 2.** Network parameters.

| Layer name | Parameter | Output shape |
| --- | --- | --- |
| Temporal convolution | filters = 100<br>kernel size = $(12, K)$<br>strides = $(2, 1)$<br>dropout = 0.1 | $(B, N, 100)$ |
| Transformer decoder | $X = 8$<br>attention head = 10<br>hidden units = 100<br>dropout = 0.5 | $(B, N, 100)$ |
| BLSTM decoder | $X = 4$<br>forward units = 120<br>hidden units = 100<br>dropout = 0.5 | $(B, N, 100)$ |
| Feed-forward 1 | units = 80 | $(B, N, 80)$ |

**Figure 3.**
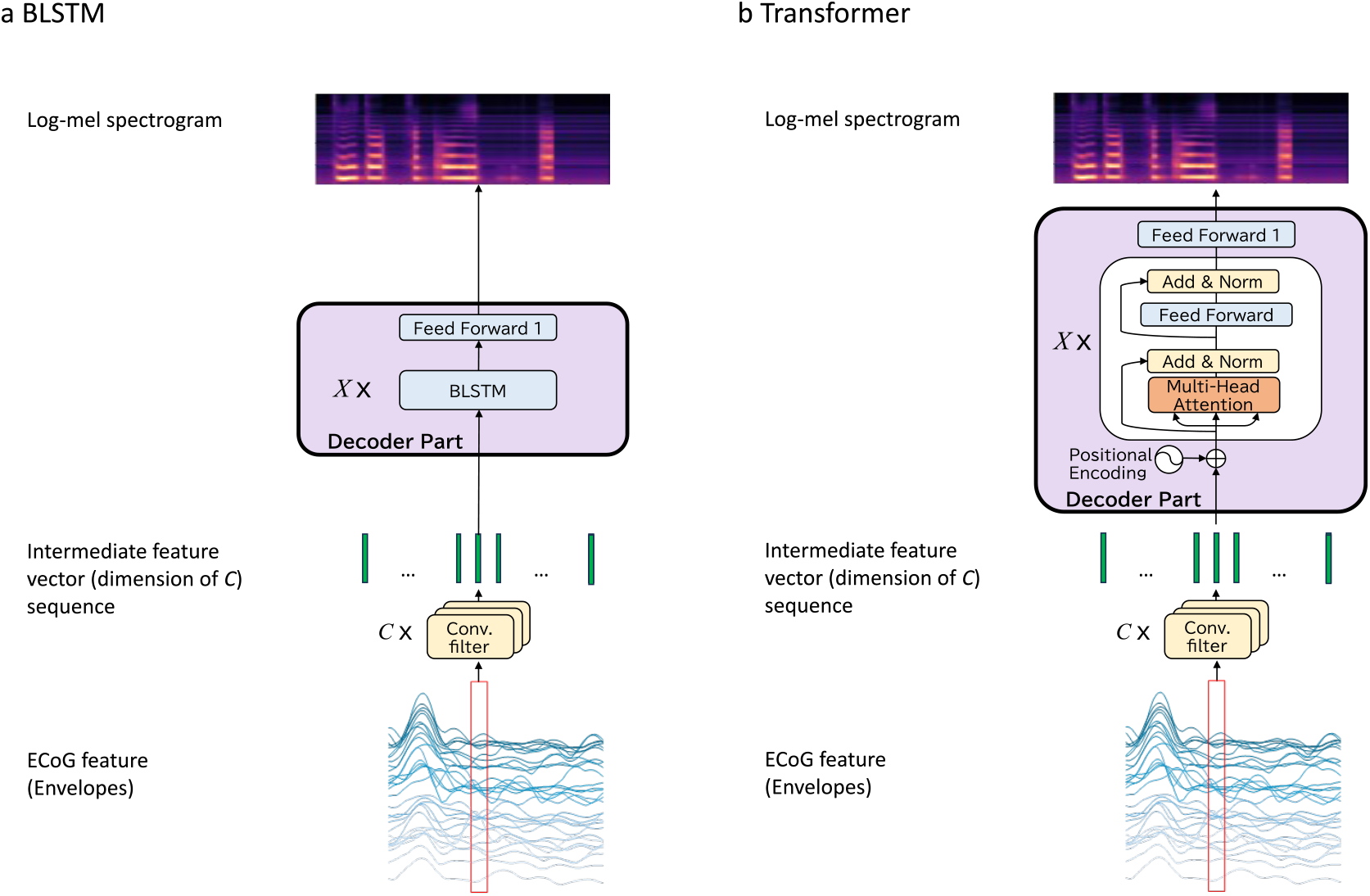
Detailed decoder architectures.

In the first stage, a temporal convolutional layer (CNN) is applied to downsample the ECoG feature sequences efficiently. The CNN input consists of *K* features, each of length *L*, where *K* is the number of electrodes (ranging from 24 to 72 across participants) and *L* ≈ 700 corresponds to a 3.5-second sequence sampled at 200 Hz. Convolutional filters with a kernel size of *W* × *K* are applied with a stride of *W*, producing an output sequence of length *N* = ⌈*L/W* ? with *C*-dimensional feature vectors.

In the second stage, the BLSTM or Transformer decoder takes the temporal convolution output as input. The decoder is composed of a stack of *X* identical layers. For the Transformer, each layer contains two sub-layers: a multi-head self-attention mechanism followed by a fully connected feed-forward network. Each sub-layer includes residual connections and layer normalization. The decoder outputs a sequence of *N* feature vectors of dimension *C*, matching the length of the input sequence. Finally, the output of the decoder stack is passed through a feed-forward network (labeled as “feed-forward 1” in Fig. 3) that transforms the feature vectors into an 80-dimensional log-mel spectrogram.

### 2.10 Neural Vocoder

In this study, we employed Parallel WaveGAN [24] § to synthesize speech waveforms. Parallel WaveGAN is a waveform generator based on a non-autoregressive WaveNet architecture and is trained to generate speech from input log-mel spectrograms. We utilized a pre-trained vocoder model developed on the JSUT corpus dataset, which contains Japanese speech recordings. Specifically, we used the model weights from “jsut parallel wavegan.v1,” which are publicly distributed online.

### 2.11. Network Implementation and Parameters

The network hyperparameters are summarized in Table 2. In the temporal convolution layer, the number of convolutional filters *C* and kernel size *W* were set to 100 and 12, respectively. In contrast, the number of electrodes *K* remained variable depending on the participants. The filters were applied with a stride size of two. The output shape variables *B* and *N* represent the batch size for training and sequence length, which depend on the data. In the decoder, we set the number of identical layers *X* to eight for the Transformer and four for the BLSTM. The number of model parameters of the Transformer and BLSTM decoders was comparable: 1, 129, 305 for the Transformer and 1, 182, 905 for the BLSTM. In “feed-forward 1,” the output dimension was set to 80, equivalent to the log-mel spectrogram dimension.

### 2.12. Network Training

Our neural networks were trained by minimizing the mean squared error (MSE) between the predicted and target log-mel spectrograms output by the decoder. The training hyperparameters are listed in Table 3. We evaluated models trained for 800, 1, 200, 1, 600, and 2, 400 epochs for both the Transformer and BLSTM architectures based on the MSE loss on the training set to determine the optimal number of training epochs. Based on this evaluation, 2, 400 epochs were selected for the Transformer and 800 epochs for the BLSTM.

**Table 3.** Training parameters.

| Parameter name | Value |
| --- | --- |
| Learning rate | 0.0005 |
| Batch size | 16 |
| Training epochs (Transformer decoder) | 2,400 |
| Training epochs (BLSTM decoder) | 800 |

For the overt speech decoding task, we used the audio signals recorded simultaneously with the ECoG signals as the ground truth. Conversely, for the covert speech decoding task, we utilized the participant’s own utterance during the overt task as the target ground truth, based on the hypothesis that the decoding framework for overt speech can be generalized to covert speech. Critically, there is no direct correspondence between covert and overt trials. Mapping variable acoustic targets during the overt task to covert neural activity, which also varies across trials, would introduce inconsistency into the training process. To address this issue and stabilize learning, we adopted a template-based approach in which a single representative utterance during the overt task was assigned as the ground truth for all covert trials of the same sentence pattern. Specifically, the audio recording from the first trial of each of the eight sentence patterns was selected as the canonical reference target. Consequently, although there were 80 trials in total, the target signals for the covert task consisted of only eight unique waveform patterns.

A key distinction in the training process between the overt and covert tasks is the variability of the ground truth signals. We employed five-fold cross-validation, with up to 80 trials per participant split into training (up to 64) and evaluation (up to 16) sets. For the overt task, since natural speech inherently contains trial-to-trial acoustic variability (e.g., fluctuations in pitch and timing) even for the same sentence, each trial constitutes a unique acoustic variation. Consequently, up to 64 distinct signal patterns were used as ground truth for training. Conversely, for the covert task, due to the use of fixed template utterances for each sentence type, only eight unique signal patterns were used as ground truth across the 64 training trials.

### 2.13. Evaluation Method

To systematically evaluate our hypothesis, we employed multiple complementary assessment methods: (1) technical quality metrics, DTW-aligned PCC; (2) analysis with Gaussian noise input to investigate the decoder’s intrinsic properties; and (3) human listener dictation tests to assess the real-world intelligibility of the synthesized speech.

The synthesized speech was evaluated using five-fold cross-validation. For all 13 participants who completed both overt and covert tasks, up to 80 trials were divided into five subsets: four subsets (up to 64 trials) were used to train the neural network model, and the remaining subset (up to 16 trials) was used for decoding the speech waveform. Separate models were trained for each participant and fold. As a result, we obtained up to 80 synthesized speech samples per participant for both overt and covert tasks for evaluation.

#### 2.13.1. Quality Evaluation

To quantitatively evaluate the quality of the reconstructed speech, we employed the DTW-aligned PCC between the log-Mel spectrograms of the original and reconstructed audio. Given the inherent temporal variability in speech production, we first applied DTW to align the two spectrograms, following the methodology described in Angrick et al. [19] and Kohler et al. [27]. Specifically, our implementation of the DTW-aligned PCC was based on the open-source code provided by Angrick et al. ∥[19] to ensure consistency with prior benchmarking studies. The alignment was performed using the FastDTW algorithm with Euclidean distance to find the optimal warping path. Let **S**_*orig*_ ∈ ℝ^*T ×F*^ and **S**_*recon*_ ∈ ℝ^*T×F*^denote the aligned log-Mel spectrograms of the original and reconstructed speech, respectively, where *T* is the number of time frames and *F* is the number of frequency bins. In our evaluation, we explicitly restricted the frequency range to the first 65 bins (*F* = 65) out of the total 80 bins. This restriction was applied because the original audio signals were recorded at a sampling rate of 9, 600 Hz, limiting the effective bandwidth to the Nyquist frequency of 4, 800 Hz. Although the Parallel WaveGAN vocoder generates audio at 24, 000 Hz, assessing frequency components beyond the original 4, 800 Hz bandwidth would be invalid. Therefore, we evaluated the correlation only within the mel-frequency bins corresponding to the valid frequency range up to approximately 4, 800 Hz. To accurately capture the spectrotemporal similarity, we calculated the DTW-aligned PCC for each frequency bin independently. For each bin *f* ∈ {1, …, *F*}, the correlation coefficient *r*_*f*_ was computed as:

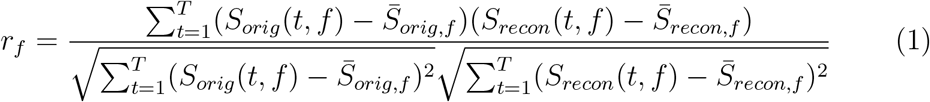

where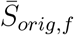and 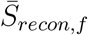represent the mean values of the *f* -th frequency bin over time. The final PCC score for each trial was defined as the average of the correlation coefficients across all frequency bins:

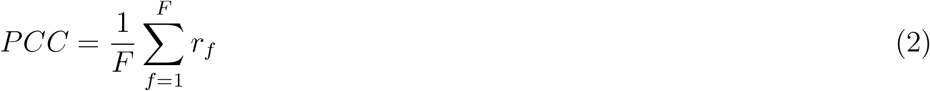

#### 2.13.2. Chance Level Estimation via Gaussian Noise Input

To rigorously assess whether the decoder truly extracts meaningful features from the neural signals rather than merely learning the statistical distribution of the target speech (i.e., overfitting to the output prior), we established a chance-level baseline using Gaussian noise as input. This procedure follows the validation strategy recommended to verify that the model performance is not driven by chance. We trained the Transformer decoder using pairs of Gaussian noise inputs and the corresponding target log-mel spectrograms. Crucially, to ensure a fair comparison with the main decoding task, the experimental conditions were kept identical to the overt/covert decoding tasks, with the sole exception of the input data. Specifically:

- **Input:** For each participant, we generated Gaussian noise signals with a duration of 3.5 seconds and a sampling rate of 9, 600 Hz. The number of channels was set to match the exact electrode count of each individual participant (median: 54 channels), ensuring the input dimensions were identical to those of the recorded ECoG signals.
- **Target:** The target log-mel spectrograms were the same representative overt speech samples used in the main decoding analysis (as described in Section 2.12).
- **Training & Evaluation:** We followed the same five-fold cross-validation protocol. The model was trained to map the Gaussian noise to the target speech, and the resulting synthesized speech from the test set was evaluated against the ground truth.

By comparing the performance of this noise-input model with that of the model trained on actual ECoG signals, we can quantify the extent to which the neural signals contribute to the decoding performance beyond the baseline statistical regularities of the speech data. Furthermore, to investigate the internal mechanism of the decoder under noise input conditions, we conducted an additional analysis by modifying the input signal length and shifting the positional encoding offsets, as detailed in Section 3.3.

#### 2.13.3. Dictation Test

To assess the real-world intelligibility of the synthesized speech, we recruited 16 listeners (see Section 2.2 for a detailed description). For each speech sample, listeners were instructed to select one sentence from a set of nine options, which included eight predefined sentences and a “not audible” option to be chosen if the sample was unrecognizable. As both the selected options (excluding “not audible”) and the reference sentences consisted of three tokens (each corresponding to one of the three phrases described in Section 2.5), we employed the TER to evaluate the accuracy of the selected sentences, defined as follows:

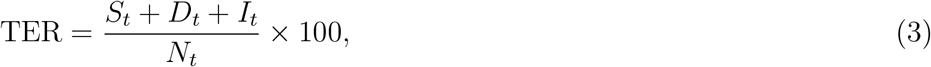

where *S*_*t*_ is the number of substituted tokens, *D*_*t*_ is the number of deleted tokens, *I*_*t*_ is the number of inserted tokens, and *N*_*t*_ is the total number of tokens in the reference sentence.

For the dictation test in the overt task, listeners were presented with anonymized synthesized speech generated by the decoder. To safeguard the privacy of the 13 participants, the audio signals used for training were voice-converted. Voice conversion was conducted using “FreeVC” [40], with the speech from the perception task employed as the target speaker for the conversion.

### 2.14. Group-level Electrode Contributions

We introduced the saliency map method [41] to determine electrode contributions to the prediction of log-mel spectrograms, following its prior application in text decoding studies [15] [31]. The saliency map is computed as the derivative of the output log-mel spectrogram with respect to the input ECoG features. Its dimensions match those of the input, with each element representing the magnitude of the contribution at a specific time point and electrode. We first calculated the variance of the saliency values along the time axis for each trial. Rather than focusing on absolute contribution magnitudes, we evaluated their relative values among electrodes by converting the electrode-wise variances to *z*-scores for each trial.

To identify contributing electrodes, we performed a statistical evaluation at the individual level. For each participant, we conducted a one-sample *t*-test (right-tailed, *µ >* 0) on the *z*-scored saliency values of each electrode across the decoding trials (approximately 80 trials). The *p*-values were adjusted using the Holm-Bonferroni method for each participant to control for multiple comparisons. Electrodes showing significantly positive activity (adjusted *p <* 0.05) were identified as “contributing electrodes.”

Subsequently, to visualize the group-level trends (*N* = 13), we aggregated the contributing electrodes from all participants onto the standard FreeSurfer pial surface image, “fsaverage” [42, 43, 44], using electrode coordinates transformed into Talairach space. The contribution magnitude for each electrode was defined as the average *z*-score across the trials. Finally, to identify the brain regions consistently recruited across both conditions, we calculated the intersection of the significant areas from the overt and covert speech tasks. A vertex was defined as a shared contributor only if it exceeded the significance threshold in both conditions.

## 3. Results

We obtained decoded log-mel spectrograms and synthesized audio signals from both overt and covert speech of 13 participants using the decoding process depicted in Fig. 1c, which includes BLSTM-based and Transformer-based decoders. Examples of decoded log-mel spectrograms and synthesized audio signals, along with the ground truth, are shown in Fig. 4 and Fig. 5 for overt speech and covert tasks, respectively. In both overt and covert speech, the log-mel spectrograms generated by the BLSTM and Transformer exhibit formant-like stripe patterns, which are characteristic of natural human speech. The Transformer notably produced temporal patterns—particularly speech onset and offset points—that more closely resembled the ground truth compared to the BLSTM outputs.

**Figure 4.**
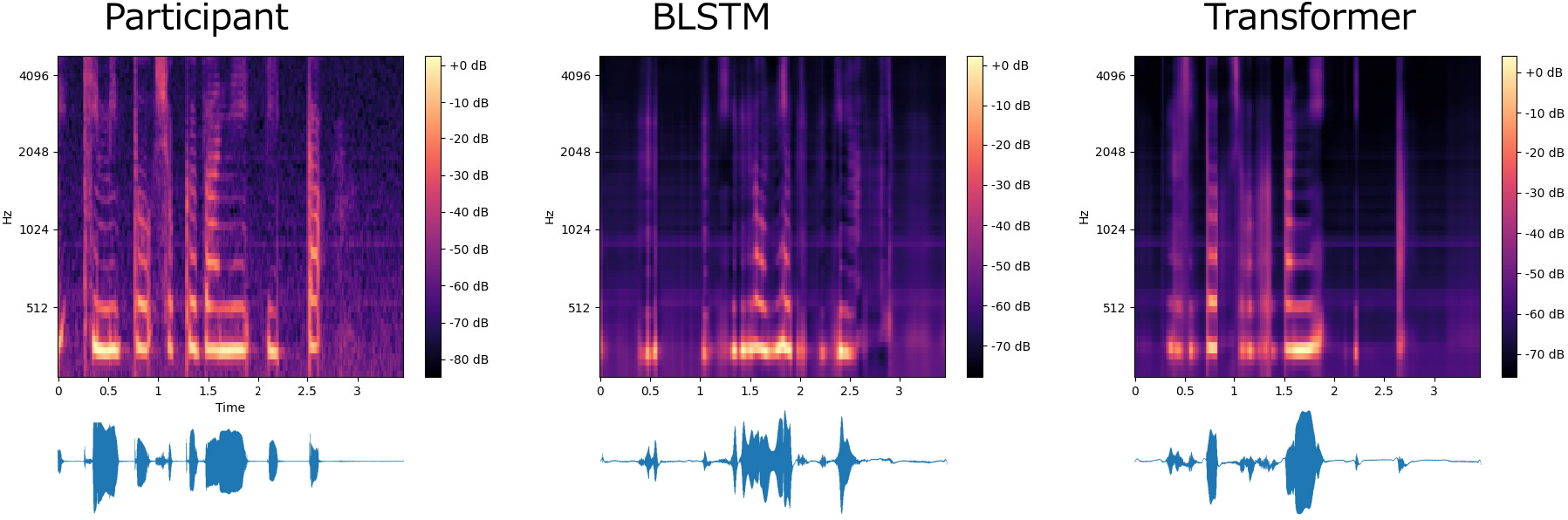
Decoded log-mel spectrograms and synthesized audio signals from overt speech: Comparison of participant (ground truth), BLSTM, and Transformer

**Figure 5.**
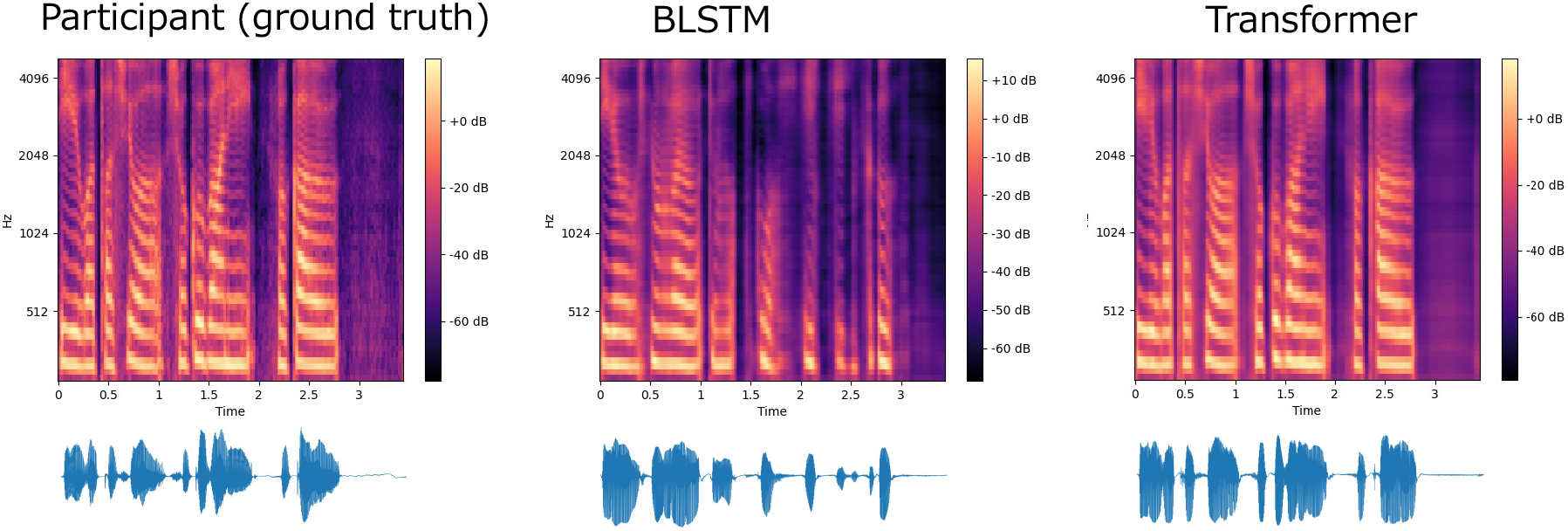
Decoded log-mel spectrograms and synthesized audio signals from covert speech: Comparison of participant (ground truth), BLSTM, and Transformer

### 3.1 Quality of Synthesized Speech

Figure 6 displays the DTW-aligned PCCs, which serve as a metric for evaluating the spectral quality of the synthesized speech for both overt (left) and covert (right) conditions. The green boxes represent the results of the BLSTM decoder, the yellow boxes represent the Transformer-based decoder, and the purple boxes represent the control condition where the Transformer model was trained and tested using Gaussian noise input (labeled “Transformer-Gaussian”). Each dot represents the average DTW-aligned PCC for an individual participant, with higher values indicating higher quality. As shown in Fig. 6, for the overt speech task, the mean DTW-aligned PCCs (mean ± s.d.) were 0.66±0.05 for the BLSTM, 0.77±0.03 for the Transformer, and 0.76±0.03 for the Transformer-Gaussian.” For the covert speech task, the mean DTW-aligned PCCs were 0.64 ± 0.06, 0.80 ± 0.03, and 0.79 ± 0.03, respectively. The Transformer model demonstrated significantly higher performance compared to both the BLSTM and the “Transformer-Gaussian” baseline across both tasks. Statistical analysis using a one-sided Wilcoxon signed-rank test with Holm-Bonferroni correction revealed significant differences between the Transformer and BLSTM (*p <* 0.001 for both overt and covert speech). Comparisons between the Transformer and “Transformer-Gaussian” were also significant (*p <* 0.001 for overt; *p <* 0.05 for covert). In terms of effect size, Cohen’s *d* values for the comparison between Transformer and BLSTM were large (*d* = 2.62 for overt, *d* = 3.38 for covert), while those between Transformer and “Transformer-Gaussian” were smaller (*d* = 0.46 for overt, *d* = 0.23 for covert).

**Figure 6.**
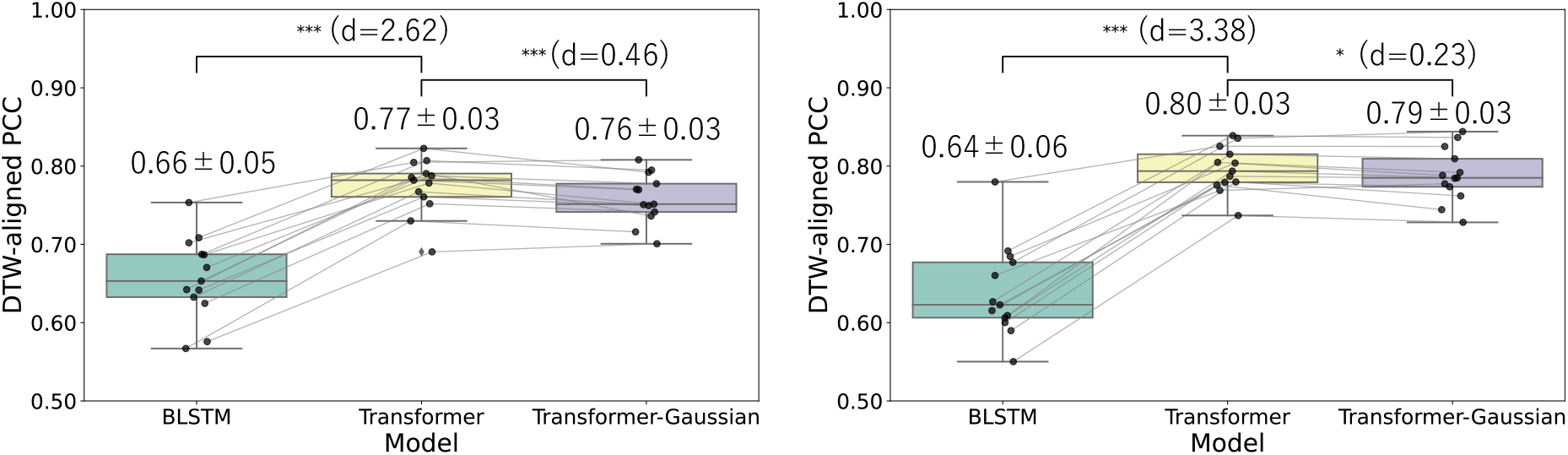
DTW-aligned PCC comparison between BLSTM and Transformer-based decoders. Box plots compare the DTW-aligned PCCs between the BLSTM and Transformer-based decoders for overt (left) and covert (right) speech synthesis tasks. Each dot represents the average DTW-aligned PCC for an individual participant, with lines connecting corresponding values across the two models. Statistical significance was assessed using a one-sided Wilcoxon signed-rank test, with the resulting *p*-values reported accordingly (* : *p <* 0.05, ** : *p <* 0.01, * * * : *p <* 0.001). The *p*-values were adjusted using the Holm-Bonferroni method.

### 3.2 Dictation Test

In the dictation test for the Transformer decoder, the “not audible” option was selected in 7.4% of trials for the overt task and 2.2% for the covert task. Notably, the selection rate for the covert task was approximately three times lower than that for the overt task.

In the subsequent results of the dictation test, the TERs for synthesized speech from the Transformer decoder, calculated based on the selected options excluding “not audible,” are presented in Fig. 7. The yellow boxes represent the Transformer-based decoder, and the purple boxes represent the control condition, where the Transformer model was trained and tested using Gaussian noise input (also labeled as “Transformer-Gaussian”). Each dot represents the average TER for an individual participant, with lower values indicating higher quality. As shown in Fig. 7, for the overt speech task, the mean TERs (mean ± s.d.) were 37.1% ± 9.0 for the Transformer and 50.8% ± 3.9 for the Transformer-Gaussian.” For the covert speech task, the mean TERs were 47.2% ± 3.9 and 51.0% ± 3.4, respectively. The Transformer model demonstrated significantly higher performance compared to the “Transformer-Gaussian” baseline across both tasks. Statistical analysis using a one-sided Wilcoxon signed-rank test with Holm-Bonferroni correction revealed significant differences between the Transformer and “Transformer-Gaussian” (*p <* 0.01 for both overt and covert speech). In terms of effect size, Cohen’s *d* values for the comparison between Transformer and “Transformer-Gaussian” were large (*d* = 1.96 for overt, *d* = 1.03 for covert).

**Figure 7.**
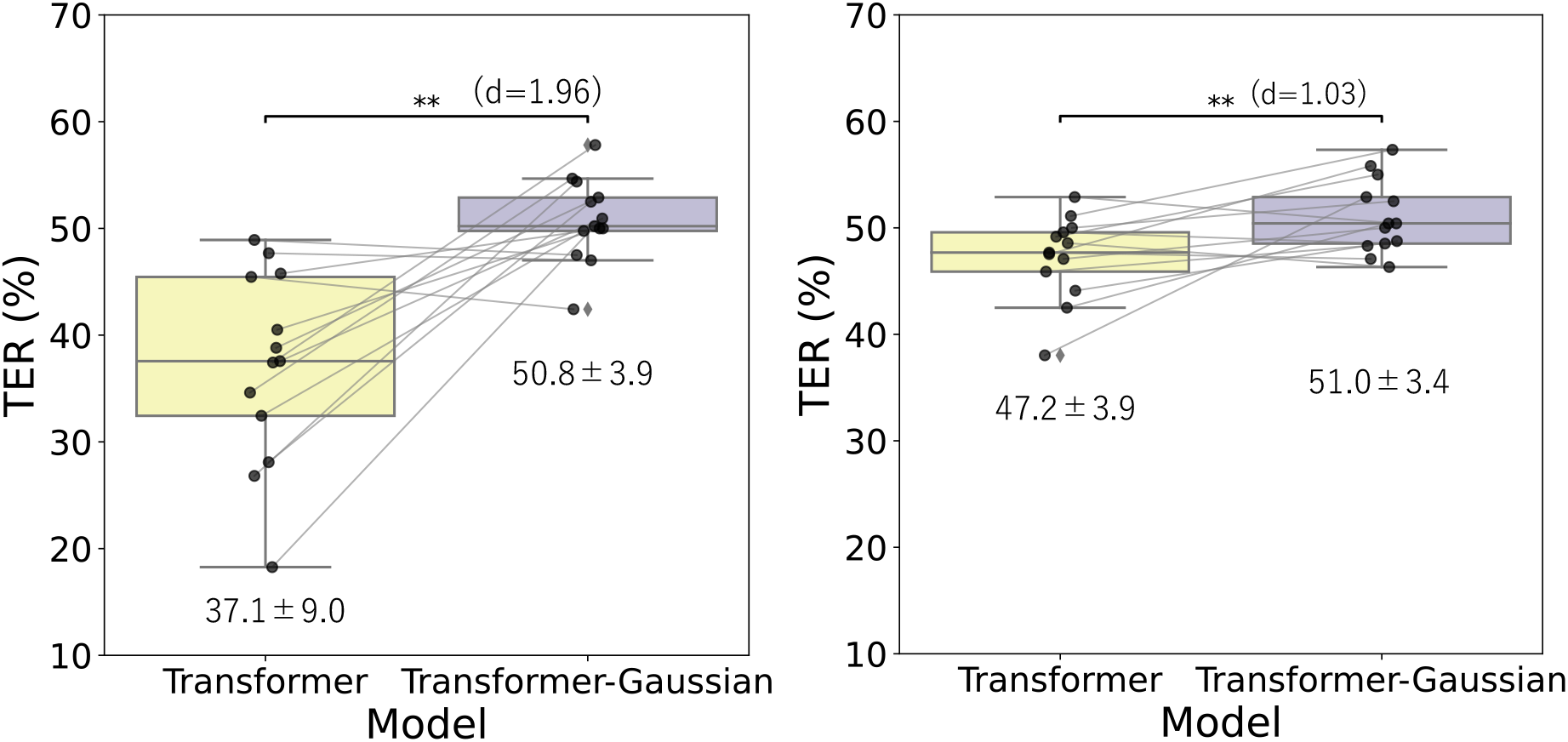
TERs by Dictation. Box plots comparing the TERs between overt and covert speech synthesis tasks. Each dot represents the average TER for one participant, and lines connect corresponding values for each participant across both models. Statistical significance was assessed using a one-sided Wilcoxon signed-rank test, with the resulting *p*-values reported accordingly (* : *p <* 0.05, ** : *p <* 0.01, *** : *p <* 0.001). The *p*-values were adjusted using the Holm-Bonferroni method.

### 3.3. Validation of Input Dependency via Gaussian Noise

Following the additional analysis procedure described in Section 2.13.2, we examined the decoder’s response to Gaussian noise inputs with modified lengths and positional encoding offsets. Figure 8 presents the log-mel spectrograms generated under these conditions. When a half-length Gaussian noise sequence was input with a positional encoding offset of 0, spectral patterns appeared corresponding to the first half of the output duration. However, as the offset was systematically adjusted (e.g., to 1*/*4 or 1*/*2 of the total length), the temporal positions of the generated spectral structures (such as formant-like patterns) shifted in alignment with the positional encoding indices. This observation demonstrates a direct correspondence between the positional encoding values and the temporal location of the generated acoustic features in the absence of meaningful input signals.

**Figure 8.**
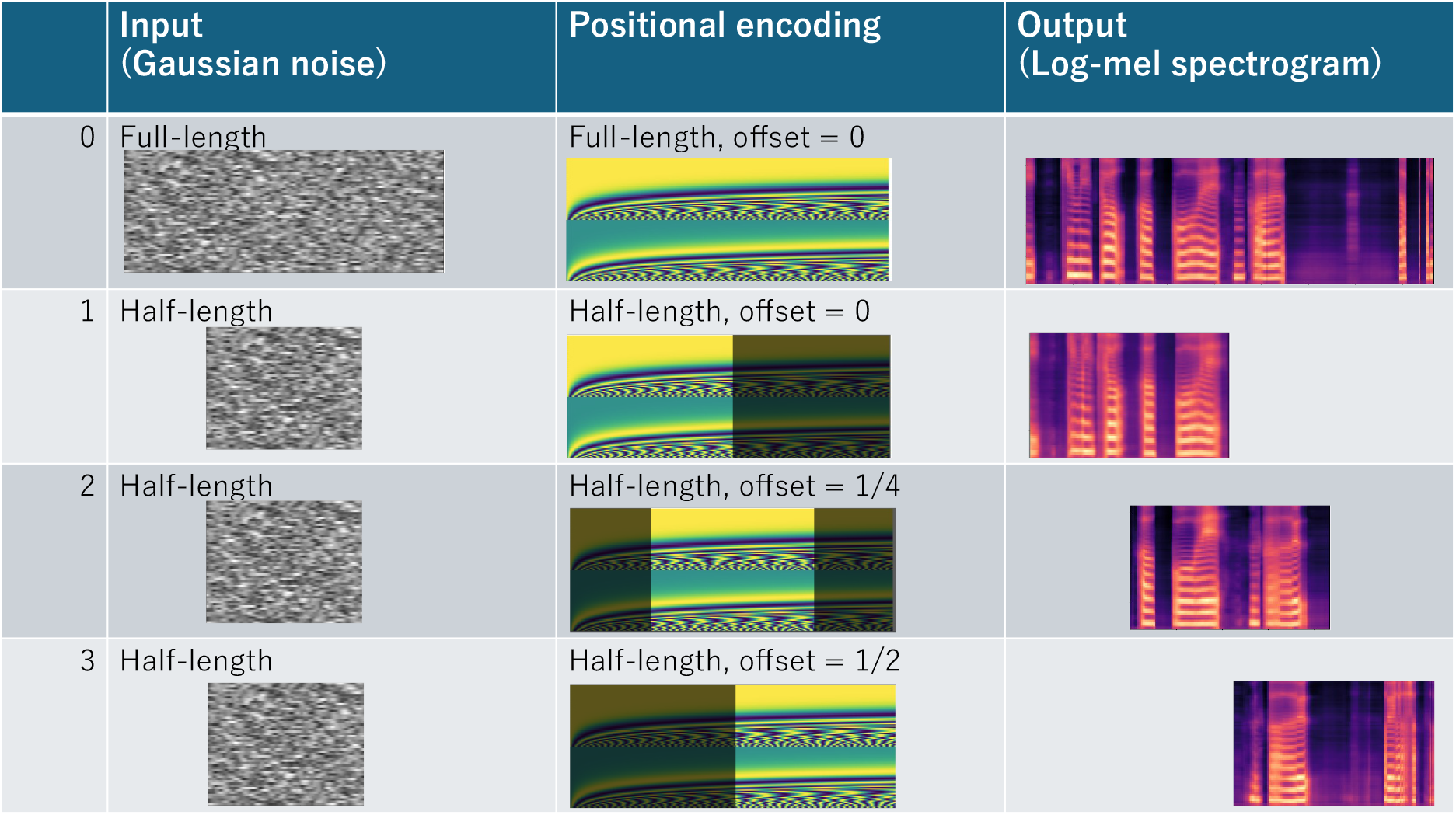
Offset control of positional encoding. Comparison of log-mel spectrogram outputs for Gaussian noise inputs with varying input lengths and positional encoding offsets applied to the Transformer-based decoder.

### 3.4 Group-level Electrode Contributions

Figure 9 illustrates the group-level electrode contribution maps (*N* = 13) on the standardized brain surface. Figure 9(a) displays the continuous contribution maps, where the left and right panels correspond to the overt and covert speech conditions, respectively. In the overt condition (Fig. 9(a), left), we observed widespread significant contributions distributed across the sensorimotor cortex, frontal lobe, temporal lobe, occipital lobe, and superior parietal lobule (SPL). A similar spatial distribution pattern was observed in the covert condition (Fig. 9(a), right).

**Figure 9.**
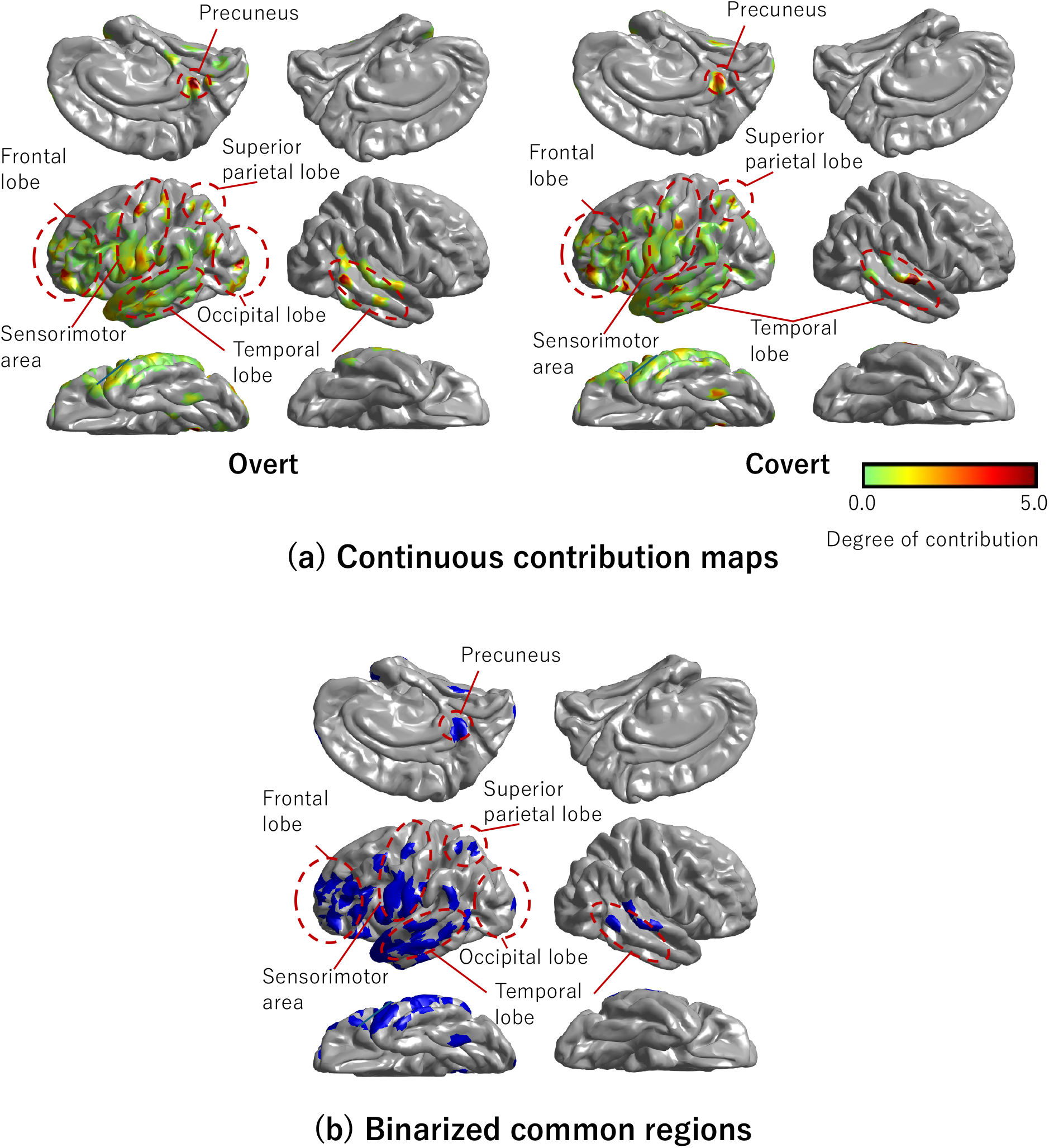
Visualization of group-level electrode contributions (*N* = 13). (a) Continuous contribution maps. The spatial distribution of contribution weights is shown for overt (left) and covert (right) conditions. Only electrodes showing statistically significant positive contributions (one-sample *t*-test, Holm-adjusted *p <* 0.05) are displayed. Warmer colors indicate higher contribution magnitudes. (b) Binarized common regions. The blue areas indicate the spatial intersection (logical AND) of the significant regions, representing vertices that exceeded the significance threshold in both the overt and covert tasks. These results highlight the consistent shared involvement of the frontal, temporal, sensorimotor area, and parietal cortices, including the precuneus.

Figure 9(b) presents the binarized common contribution regions. In this map, the blue areas indicate the spatial intersection of electrodes that showed significant contributions (Holm-Bonferroni-corrected *p <* 0.05) in both conditions. Visual inspection revealed that these shared neural representations were robustly located in the sensorimotor cortex, frontal lobe, temporal lobe, SPL, and precuneus.

## 4. Discussion

The primary goal of this study was to validate the hypothesis that a decoding framework trained on utterances during the overt task could be successfully generalized to synthesize covert speech. Our results strongly support this hypothesis, demonstrating that using the participant’s own utterance as a surrogate ground truth enables effective speech synthesis from covert neural activity. Building on this confirmation, we identified three key findings that warrant detailed discussion: (1) The Transformer-based decoder consistently outperformed the BLSTM in both overt and covert speech synthesis in terms of DTW-aligned PCC (*p <* 0.001). (2) Even with Gaussian noise input, the decoder generated log-mel spectrograms with high structural quality (high DTW-aligned PCC); however, the dictation test revealed that accurate semantic content generation strictly requires information from ECoG signals. (3) Analysis using saliency maps revealed patterns of neural contributions that were consistent with those identified in our previous study on text decoding, suggesting a commonality in the brain regions involved in both speech and text decoding tasks.

These findings provide important insights into the neural decoding of covert speech and highlight the potential of Transformer-based architectures for speech BCIs. In the following, we discuss the results from three aspects.

### 4.1. Comparison of Transformer and BLSTM

For the first aspect, the Transformer outperformed BLSTM in terms of DTW-aligned PCC. Previous studies by Komeiji et al. [16, 31] and Shigemi et al. [29] have demonstrated that the Transformer surpasses BLSTM for both text decoding and speech synthesis from ECoG signals. Consistent with these findings, the ability to capture long-range dependencies in the input ECoG signals appears to play a crucial role in speech synthesis. Although Kohler et al. [27] achieved a PCC of up to 0.8 using GRU, a type of RNN similar to BLSTM, for decoding Dutch sentences from sEEG, our results suggest that replacing RNN-based architectures with a Transformer can further improve result quality. Furthermore, Chen et al. [45] applied a Swin Transformer (a variant of Transformer) [46] to decode English speech from ECoG or sEEG and reported correlations exceeding 0.8. Therefore, considering the consistent superiority shown in this and prior studies, it is anticipated that Transformer architectures will become the de facto standard for speech synthesis from ECoG or sEEG.

### 4.2 Ability to Generate High-quality Log-mel Spectrograms

For the second aspect, we analyzed the remarkable ability of the Transformer decoder to generate high-quality log-mel spectrograms. Interestingly, this capability was observed even when Gaussian noise was used as input (Fig. 6). Further analysis revealed that positional encoding plays a crucial role in producing this high-quality output by providing temporal context to the input (Fig. 8). Even when Gaussian noise, which inherently lacks meaningful context, is used as input, the positional encoding enables the Transformer’s output to maintain a speech-like temporal structure. This suggests that the Transformer decoder functions effectively as a pattern generator, capable of reproducing the learned statistical structure of human speech somewhat independently of the specific input characteristics.

This behavior parallels recent advancements in generative AI, such as Stable Diffusion [47], where neural networks demonstrate the capacity to generate high-fidelity images from noise vectors. In the case of Stable Diffusion, the model–trained on massive datasets like LAION-400M [48]–learns the underlying distribution of image data, allowing it to synthesize coherent structures from random noise. Similarly, our results imply that the Transformer decoder has learned the statistical priors of the target speech spectrograms. This characteristic explains why the synthesized speech from Gaussian noise achieved high DTW-aligned PCC; the model successfully reconstructed the “texture” of speech. Stable diffusion technology has also been extended to estimate images from fMRI data [49], further highlighting the relevance of generative capabilities in neural decoding.

This generative capability also elucidates the apparent discrepancy observed between overt and covert speech results. Specifically, synthesized covert speech exhibited higher DTW-aligned PCCs (0.80 ± 0.03) compared to overt speech (0.77 ± 0.03, Fig. 6), along with a lower proportion of “not audible” responses in the dictation test (see Section 3.2). These metrics might superficially imply that the acoustic quality of synthesized overt speech is inferior to that of covert speech, despite covert speech demonstrating inferior semantic accuracy (TER). This phenomenon can be attributed to the difference in target variability during training. For covert speech, we utilized a single representative overt speech sample as a fixed template target for all trials within the same sentence pattern (as described in Methods). Consequently, the model converged to this single, clean spectral pattern, resulting in high structural similarity (high DTW-aligned PCC). In contrast, for overt speech, the model was trained on the actual vocalizations from each trial, which inherently contain acoustic variability (e.g., pitch and timing fluctuations). Reproducing such trial-by-trial variability is a more complex task, naturally yielding slightly lower DTW-aligned PCCs compared to the fixed-target condition.

Crucially, the dictation test results (TER) clarify that high spectral similarity does not equate to semantic accuracy. While the covert task yielded high DTW-aligned PCCs due to the low-variance target, the overt task achieved better TER (37.1% vs 47.2%), reflecting the richer and more reliable neural representations available during actual vocalization. Furthermore, the noise input condition achieved a high DTW-aligned PCC but poor semantic accuracy (chance-level TER). In other words, the Transformer demonstrated the capability to extract meaningful features from ECoG signals to construct semantically accurate synthesized speech, distinguishing true decoding from mere pattern generation.

These findings lead to an important conclusion regarding the future direction of speech BCI. The challenge of synthesizing high-fidelity, natural-sounding speech (high DTW-aligned PCC) has been largely overcome by combining the Transformer decoder and a pre-trained neural vocoder; the model can autonomously generate the necessary acoustic fine structure. Therefore, the primary focus of future research should shift from merely pursuing signal quality to improving the extraction of precise semantic content from neural activity. Improving the decoding of “content” will likely require larger training datasets to better map neural patterns to linguistic representations, rather than just acoustic textures.

### 4.3 Electrode Contributions

Regarding the third aspect, our previous study [31] demonstrated that a model trained on overt speech could be directly applied to decode text from covert speech. In this study, we investigated the anatomical basis of this transferability to determine whether a similar cross-modal approach is feasible for speech synthesis from ECoG signals. Establishing this transferability is crucial because it allows models to be trained on overt speech–where the speaker’s own actual speech serves as a reliable ground truth label– thereby eliminating the need for complex experimental paradigms like the “Karaoke-like text highlighting” typically required to collect labeled covert speech.

To this end, we examined the brain regions that commonly contribute to speech synthesis in both overt and covert conditions. As shown in Figure 9(C), we identified robust common contributions in four key anatomical areas: the frontal lobe, temporal lobe, parietal lobe, and precuneus.

In the frontal region, specifically the rostral lateral prefrontal cortex (LPFC), we observed significant common activity. This region is known to be activated, with left-hemisphere dominance, during task-set maintenance and verbal working memory tasks [50], suggesting its role in holding the intended sentence in working memory. In the temporal lobe, the shared activity likely reflects the processing of auditory feedback during overt speech and the prediction of auditory imagery during covert speech, consistent with the dual-stream model of speech processing [51].

Furthermore, the activation in the SPL is particularly noteworthy. This region, including its medial component (i.e., the precuneus), acts as a central node for visuospatial imagery, episodic memory retrieval, and internal mentation [52]. While the inferior parietal cortex is often linked to the “intention to speak” [53], the SPL plays a complementary role in sensory-motor integration and the maintenance of internal representations needed for motor planning. Therefore, the shared activity in the SPL and the precuneus suggests that both overt and covert speech tasks involve the generation and maintenance of internal imagery (visual or kinesthetic) related to the text prompts, as well as the internal monitoring of speech production.

Finally, the significant shared contribution in the sensorimotor cortex aligns with the simulation theory of action [54]. This theory posits that covert actions (such as covert speech) recruit the same neural mechanisms involved in programming and preparing actual movements, with inhibition occurring only at the executive output level. This implies that the Transformer decoder leverages these shared motor-planning signals, which persist even in the absence of overt vocalization, to reconstruct speech features.

Importantly, the finding that high-level cognitive networks—encompassing planning, auditory imagery, and memory—are consistently activated across both conditions provides a strong neurophysiological rationale for our approach. These shared neural representations suggest that a model trained on overt speech learns to capture the underlying intent or imagery of speech rather than just motor commands. This result strongly supports the feasibility and motivation for future work focused on applying overt-trained models to covert speech synthesis.

## 5. Conclusion

This study demonstrated the successful synthesis of speech from neural activity during covert reading using ECoG signals. By employing a Transformer-based decoder in combination with a pre-trained neural vocoder, Parallel WaveGAN, we were able to generate high-quality audio despite the limited size of the training. Experiments involving ECoG signals from 13 participants yielded synthesized speech with DTW-aligned PCC ranging from 0.74 to 0.84. These results highlight the effectiveness of the Transformer decoder in accurately reconstructing high-fidelity log-mel spectrograms from ECoG signals, even under data-constrained conditions. The ability to synthesize intelligible speech from covert reading underscores the potential of this approach for advanced speech BCI applications, particularly for enabling communication in individuals with speech impairments.

## Acknowledgment

This work was partly supported by JSPS KAKENHI, 20H00235, and 23H00548.

## Footnotes

‡ The source code for the network architecture and training procedures is publicly available at https://github.com/ttlabtuat/ECoG2Speech.

§ Implemented as available at https://github.com/kan-bayashi/ParallelWaveGAN

∥ Available at https://github.com/cognitive-systems-lab/closed-loop-seeg-speech-synthesis/blob/master/eval_steps/exp2.py

